# Does *Borrelia burgdorferi* sensu lato facilitate the colonisation of marginal habitats by *Ixodes ricinus*? A correlative study in the Swiss Alps

**DOI:** 10.1101/273490

**Authors:** Mélissa Lemoine, Luca Cornetti, Barbara Tschirren

## Abstract

Parasites can alter host and vector phenotype and thereby affect ecological processes in natural populations. Laboratory studies have shown that *Borrelia burgdorferi* sensu lato, the causative agent of human Lyme borreliosis, induces physiological alterations in its main tick vector in Europe, *Ixodes ricinus*, which increase its survival under challenging conditions. We hypothesise that these phenotypic alterations may allow *I. ricinus* to colonise marginal habitats, thereby fuelling the ongoing range expansion of *I. ricinus* towards higher elevations and latitudes induced by climate change. To explore the potential for such an effect under natural conditions, we studied the prevalence of *B. burgdorferi* s.l. in questing *I. ricinus* and its variation with elevation in the Swiss Alps. We screened for *B. burgdorferi* s.l. infection in questing nymphs of *I. ricinus* (N = 411) from 15 sites between 528 and 1774 m.a.s.l to test if *B. burgdorferi* s.l. prevalence is higher at high elevations (i.e. in marginal habitats). We found that *B. burgdorferi* s.l. prevalence in *I. ricinus* nymphs decreased linearly with increasing elevation and that it was 12.6% lower in *I. ricinus* nymphs collected at high elevations compared to nymphs in the core range. Thus, we found no evidence that the *B. burgdorferi* s.l.-induced alterations of *I. ricinus* phenotype facilitate the colonisation of marginal habitats in the wild. These findings have implications for a better understanding of eco-evolutionary processes in natural host-parasite systems, as well as the assessment of Lyme borreliosis risk in regions where *I. ricinus* is newly emerging.

## 1. Introduction

Parasites can alter the physiology, morphology and behaviour of their hosts (Poulin, 1994; Price, 1980; Thomas et al., 2002). Vectors can similarly be the target of parasite manipulation (Dobson, 1988; Hurd, 2003). For example, *Yersinia pestis* induces aggressive feeding in fleas and the regurgitation of the pathogen during feeding (Bacot and Martin, 1914). All these manipulations increase the chances of parasite transmission, and are thus adaptive from the parasite’s perspective (Lefevre and Thomas, 2008). However, not all phenotypic alternations are the result of such adaptive manipulations (Cezilly et al., 2013; Thomas et al., 2005). Phenotypic changes in infected hosts and vectors can also be a side effect of infection that is of no adaptive value for the parasite.

Surprisingly little is known about the ecological consequences of parasite-induced phenotypic alterations in hosts or vectors. Recent studies have suggested that parasite-induced changes in physiology, morphology or behaviour may have profound effects on host population dynamics, trophic niche specialisation as well as interactions between competitors and predators at all trophic levels (Britton and Andreou, 2016; Dunn et al., 2012; Wood and Johnson, 2015). Consequently, parasites can have a strong ecological impact that becomes especially apparent when they are missing or introduced, e.g. during biological invasions (Hatcher et al., 2014; Wood and Johnson, 2015). Invasive host populations often harbour fewer parasites than populations in their native range, which may enhance their population growth and competitive ability (i.e. enemy release hypothesis, Heger and Jeschke, 2014). Parasites may be ‘lost’ because of low host densities and founder effects during the three phases of an invasion: introduction, establishment and range expansion (Kolar and Lodge, 2001). However, empirical studies investigating the role of parasites during range expansion from core populations, i.e. without bottlenecks occurring during the introduction and establishment phases, are still rare (White and Perkins, 2012), and both increasing and decreasing parasite loads at the host range margins have been reported (e.g. Briers, 2003; Patot et al., 2010; Phillips et al., 2010).

The distribution of *Ixodes spp.* ticks is strongly influenced by abiotic factors, such as temperature and humidity, and there is accumulating evidence that in Europe and North America *Ixodes spp.* ticks have been expanding northwards as well as towards higher elevations during the last decades due to climate warming (see Medlock et al., 2013; Ostfeld and Brunner, 2015 for reviews). In Europe, *I. ricinus* is the main vector of *Borrelia burgdorferi* sensu lato, the causative agent of human Lyme borreliosis. *Borrelia burgdorferi* s.l. forms a complex comprising of at least 18 genospecies (Margos et al., 2011). A diverse host community, which includes rodents, insectivores, birds and reptiles, acts as reservoir hosts for *B. burgdorferi* s.l.. In Europe, *B. afzelii* and *B. garinii* are the most common genospecies (Rauter and Hartung, 2005). *Borrelia afzelii* is a rodent specialist, infecting mice (*Apodemus sylvaticus and A. flavicollis*) and voles (*Myodes glareolus and Microtus agrestis*), but also shrews (*Sorex araneus and S. minutus;* Hellgren et al., 2011), whereas *B. garinii* is a bird specialist, infecting a range of bird species such as *Turdus* spp., *Sturnus* vulgaris, *Sylvia* spp. and *Parus major* (Comstedt et al., 2006; Margos et al., 2011). Laboratory studies have found that *B. burgdorferi* s.l.-infected *I. ricinus* are bigger, quest longer for a host, have higher energy reserves (for the same body mass) and survive better in harsh environments than uninfected *I. ricinus*, suggesting that *B. burgdorferi* s.l. induces phenotypic alterations in the vector (Herrmann and Gern, 2015, and references therein). These alterations of tick physiology and behaviour may facilitate the ongoing range expansion, and lead to particularly high prevalence of *B. burgdorferi* s.l. in *I. ricinus* populations at range margins. Although assessing the role of *B. burgdorferi* s.l. in influencing vector range expansion to marginal habitats is crucial for our understanding of eco-evolutionary processes in natural host-parasite systems, but also for quantifying public health threats in regions where ticks are newly emerging, to our knowledge no study has tested this hypothesis in *I. ricinus* populations under natural conditions to date.

Although exclusion experiments (i.e., where a taxon is excluded) are the gold method to assess the ecological role of parasites, such manipulations remain an enormous practical and logistic challenge in *I. ricinus* and its pathogens. Moreover, experiments in the wild testing whether *B. burgdorferi* s.l. enhances survival and reproduction of *I. ricinus* under high elevation conditions (i.e., physiological alterations) would only provide a partial picture because *B. burgdorferi* s.l. might affect dispersal abilities of *I. ricinus* through its ability to find a host (i.e., behavioural alterations).

Because of these practical challenges, we used a correlational approach in naturally infected tick populations in the Swiss Alps to evaluate the potential of *B. burgdorferi* s.l. to facilitate the colonisation of marginal habitats by *I. ricinus*. Previous studies monitored *B. burgdorferi* s.l. and *I. ricinu*s within their core range (Aeschlimann et al., 1987; Gern et al., 2008; James et al., 2013; Jouda et al., 2004; Moran Cadenas et al., 2007; Stünzner et al., 2006), at the range margins without a direct comparison with core populations (Ragagli et al., 2016) or when *B. burgdorferi* s.l. prevalence was low (Daniel et al., 2009; Danielová et al., 2010; Danielová et al., 2006). In our study we directly compared tick populations at different elevations, including core range and range margin populations. We predict that if *B. burgdorferi* s.l. alters the phenotype of *I. ricinus,* thereby making them better colonizer or long-distance dispersers, *B. burgdorferi* s.l. prevalence will be disproportionally high at the range margin (i.e. at high elevations). Furthermore, we predict that if *B. burgdorferi* s.l. enhances specifically *I. ricinus* survival under harsh abiotic conditions, *B. burgdorferi* s.l. prevalence and *I. ricinus* abundance will be positively correlated within the core range of *I. ricinus*.

## 2. Material and Methods

### 2.1. Study area

We sampled questing *I. ricinus* at fifteen sites across an elevational range from 528 to 1774 m.a.s.l. in the Swiss Alps (Kanton Graubünden, Table 1). Each site was visited three times, once in June, July and August 2014 to obtain an average measure of questing tick abundance per site. During each sampling session, four people dragged a white blanket (1 m x 1 m each) slowly over the ground vegetation during 15 min each. During these 15 min, each collector regularly checked for ticks on their blanket and transferred them to Eppendorf tubes containing 95% ethanol. Although an area-based estimation is often more accurate than a time-based approach, it was not possible to apply such a method in our study because of the rugged mountain terrain of the Swiss Alps. We did not observe a significant difference in questing tick abundance patterns according to elevation across the season (negative binomial distribution for count data: *θ* = 1.12; elevation x sampling session: *χ*^2^_2_ = 0.86, *P* = 0.649; elevation: *χ*^2^_1_ = 30.65, *P* <0.001; sampling session: *χ*^2^_2_ = 6.933, *P* = 0.031). Tick life stage and species were verified with a dissection microscope in the laboratory on the basis of morphologic features following Hillyard (1996).

**Table 1.** *Borrelia burgdorferi* s.l. prevalence in questing nymphs at 15 sites in the Swiss Alps. Sampling site and acronym, GPS coordinates, elevation, number of *I. ricinus* adults and nymphs collected per h during the three sampling sessions (in June, July and August), average number of ticks (adults & nymphs) dragged per hour (Tick / h) and tick range position of the site (range core (C), range margin (M) and outside of the range (O)) are given. Furthermore, the number of nymphs screened for *Borrelia* infection (analysed nymphs), the number of nymphs infected with *B. afzelii* and other *B. burgdorferi* s.l. genospecies and the infection prevalence (%) considering all *B. burgdorferi* s.l. genospecies is reported.

| Site | Label | GPS Coordinates |  | Elevation<br>(m.a.s.l.) | Adults<br>/ h | Nymphs<br>/ h | Ticks<br>/ h | Range<br>exp. | Analysed<br>nymphs | Borrelia<br>afzelii | Borrelia<br>others | % nymphs<br>infected |
| --- | --- | --- | --- | --- | --- | --- | --- | --- | --- | --- | --- | --- |
| Untevaz | UNT | 46.938 | 9.547 | 528 | 7,4,0 | 46,12,5 | 24.7 | C | 34 | 10 | 2 | 35 |
| Malans | MAL | 46.992 | 9.558 | 560 | 7,2,1 | 11,32,5 | 19.3 | C | 35 | 4 | 2 | 17 |
| Rodels | ROD | 46.761 | 9.426 | 630 | 5,8,0 | 42,11,10 | 25.3 | C | 34 | 10 | 4 | 41 |
| Sagogn | SAG | 46.783 | 9.233 | 693 | 44,5,8 | 7,7,5 | 25.3 | C | 35 | 8 | 2 | 28 |
| Passug | PAS | 46.841 | 9.538 | 732 | 29,22,5 | 98,108,35 | 99 | C | 34 | 9 | 0 | 26 |
| Trimmis | TRI | 46.882 | 9.56 | 762 | 12,12,11 | 109,63,99 | 102 | C | 35 | 6 | 4 | 28 |
| Bonaduz | BON | 46.799 | 9.353 | 944 | 14,11,8 | 38,10,20 | 33.7 | C | 32 | 6 | 0 | 18 |
| Castiel | CAS | 46.826 | 9.57 | 1094 | 6,8,3 | 50,55,11 | 44.3 | C | 34 | 4 | 2 | 17 |
| Seewis | SEE | 46.996 | 9.637 | 1106 | 2,8,3 | 38,45,19 | 35.3 | C | 35 | 8 | 0 | 22 |
| Tomils | TOM | 46.772 | 9.454 | 1138 | 2,12,4 | 7,77,4 | 38.3 | C | 34 | 15 | 1 | 47 |
| Flims | FLI | 46.827 | 9.281 | 1144 | 4,0,0 | 13,4,3 | 8 | C | 35 | 3 | 0 | 8 |
| Ruschein | RUS | 46.795 | 9.169 | 1454 | 1,0,1 | 13,9,4 | 9.3 | R | 22 | 1 | 0 | 4 |
| Praden | PRA | 46.818 | 9.59 | 1582 | 1,0,0 | 1,9,1 | 4 | R | 9 | 0 | 0 | 0 |
| Feldis | FEL | 46.789 | 9.453 | 1673 | 0,0,0 | 1,1,0 | 0.7 | R | 3 | 0 | 0 | 0 |
| Vilan | VIL | 47.020 | 9.583 | 1774 | 0,0,0 | 0,0,0 | 0 | O | 0 | 0 | 0 | 0 |

Questing *I. ricinus* abundance (i.e. the average number of *I. ricinus* nymphs and adults sampled per dragging hour across all sampling sessions) was high up to an elevation of 1100 m.a.s.l. (except for FLI), and then strongly decreased (see Table 1, Fig. 1A). No questing *I. ricinus* were found above 1673 m.a.s.l.. We therefore defined sites between 1450 and 1680 m.a.s.l. to represent the elevational range margin of *I. ricinus* ticks in our study area (note that no sites at 1150-1450 m.a.s.l. were included in our study).

**Figure 1.**
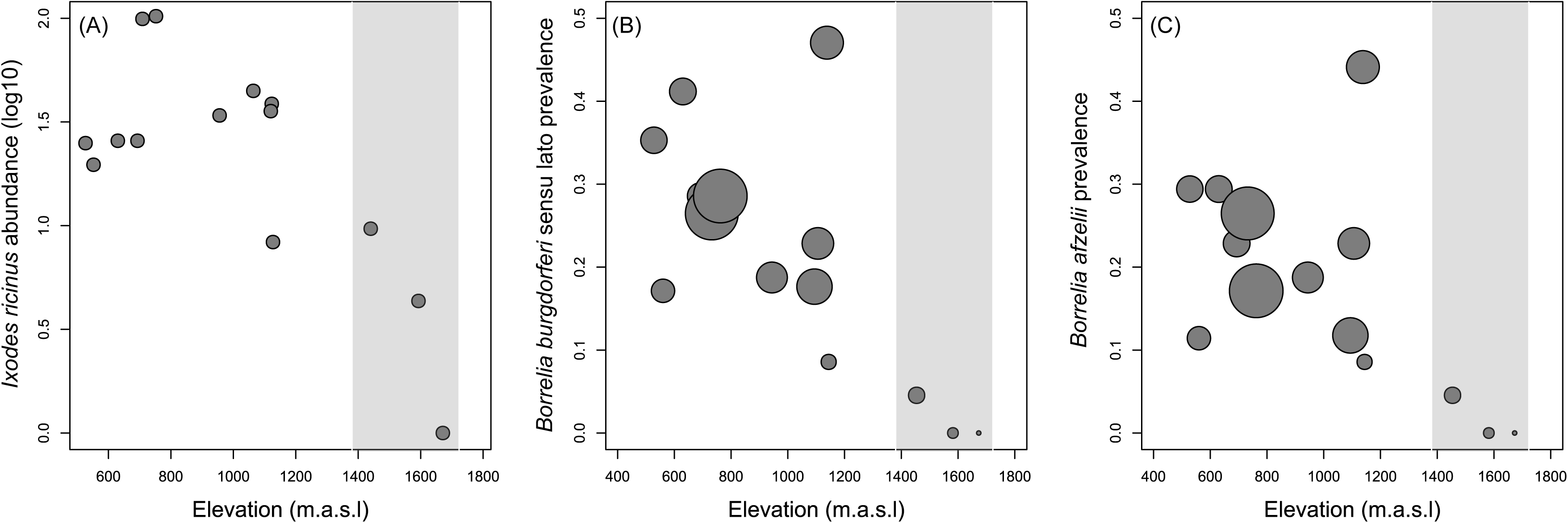
Abundance of questing *Ixodes ricinus* ticks (log(average number of nymphs and adults dragged per hour + 1)) in the vegetation (A) and prevalence of *Borrelia burgdorferi* sensu lato (B) and *Borrelia afzelii* (C) in *I. ricinus* nymphs at different elevations (m.a.s.l.) in the Swiss Alps. The grey area represents the range margin of *I. ricinus* ticks. The size of the dots in (B) and (C) is proportional to the abundance of questing *I. ricinus* nymphs.

### 2.2. *Borrelia burgdorferi* s.l. infection in questing *Ixodes ricinus* nymphs

Because questing *I. ricinus* abundance varied widely among sites (Table 1), and because we collected at least four times more nymphs than adults, we randomly selected 34-35 questing *I. ricinus* nymphs per site for *B. burgdorferi* s.l. detection (or all nymphs that were collected if < 35; Table 1). DNA was extracted using the ‘HotShot’ method (Råberg, 2012; Truett et al., 2000) with slight modifications. Each nymph was incubated with 60 μL of alkaline lysis reagent (25 mM NaOH, 0.2 mM EDTA) and a metal bead (2 mm in diameter) at 95°C for 15 min. After homogenisation with a Retsch TissueLyser (Haan, Germany) for 2 min, the samples were incubated at 95°C for 15 min. After cooling, 60 μL 40 mM Tris–HCl was added to each tube.

To assess the *B. burgdorferi* s.l. infection status of *I. ricinus* nymphs, we used a combination of two complimentary approaches. First, we specifically targeted *B. afzelii* using a highly sensitive quantitative real-time PCR (qPCR) assay using the *B. afzelii*-specific primers Fla5F 5‘-CACCAGCATCACTTTCAGGA-3’ and Fla6R 5‘-CTCCCTCACCAGCAAAAAGA-3’ (Råberg, 2012) on a StepOnePlus Real-Time PCR machine (Applied Biosystems). Each reaction contained 10 μL of SYBER Select Master Mix (Applied Biosystems), 0.8 μL of each primer (10 μM), 4.4 μL of water and 4 μL of template DNA in a final volume of 20 μL. Two series of four standards, two negative and two positive tick samples, as well as two no-template controls were included in each run. The PCR amplification protocol consisted of an initial denaturation step at 50°C and 95 °C for 2 min each, followed by 42 cycles of 95°C for 15 s, 59°C for 30 s, and 72°C for 30 s. The length of the amplicon was 129 bp and the melting temperature was between 77.3°C and 77.9°C. A 100% repeatability was obtained based on 40 samples (10 positives and 30 negatives) when amplification occurred before 34 cycles. Out of 85 *B. afzelii-*positive samples, only two samples amplified between 34 and 37 cycles. These samples were repeated and considered to be *B. afzelii*-infected only if they were found to be positive twice.

In a second step, all samples that were found to be *B. afzelii*-negative using the qPCR approach described above were screened for the presence of other *B. burgdorferi* s.l. genospecies using 16S-LD primers designed to target all *Borrelia* genospecies associated with Lyme borreliosis (Marconi and Garon, 1992). Amplifications were performed in a total volume of 10 μL containing 0.2 μL JumpStart Taq DNA Polymerase (Sigma-Aldrich), 0.5 μL of each primer (300 nM; LD-F: 5‘-ATGCACACTTGGTGTTAACTA-3’ and LD-R: 5‘-GACTTATCACCGGCAGTCTTA-3‘) and 2 μL of DNA template. The PCR amplification protocol consisted of an initial denaturation step at 94°C for 1 min, followed by 42 cycles of denaturation at 94°C for 30 s, annealing at 50°C for 30 s, and extension at 72°C for 90 s, with a final elongation step at 72°C for 10 min. PCR products were visualized under UV light on 1% agarose gels that were stained with SYBR® Safe DNA gel stain (Thermo Scientific). The length of the amplicon was 351 bp. We used DNA from seven reference genospecies (B31 and Pbre (*B. burgdorferi* s.s), Phei (*B. garinii*), Pko and PVPM (*B. afzelii*) and Pbi (*B. bavariensis*) to verify that they are detected with the 16S approach. All references were successfully amplified.

Each run included *B. afzelii*-positive samples identified with the qPCR (see above) as controls. All of these samples were found to be *B. burgdorferi* s.l.-positive using the 16S method. The repeatability based on 39 samples (12 negative and 27 positive samples) was 85 %. Out of the six samples that were not repeatable with the 16S method, three were previously quantified with the qPCR method and were found to have very low *B. afzelii* DNA concentrations (i.e. more than 30 cycles on the qPCR) suggesting that the sensitivity of the 16S approach decreases at low infection intensities. During the development of the methods, some fragments were sequenced and all were found to be *B. burgdorferi* s.l. suggesting that both methods were specific to the Lyme borreliosis causing genospecies.

### 2.3. Statistical analyses

*Borrelia* infection (i.e. *B. burgdorferi* s.l. or *B. afzelii* infection) in *I. ricinus* nymphs was analysed using a generalized mixed effect model with a binomial error structure where each nymph was treated as an individual observation (0 = non infected, 1 = infected) and site ID was included as a random effect. In a first step, we tested for an association between *B. burgdorferi* s.l. or *B. afzelii* prevalence and elevation by including elevation and elevation^2^ as fixed effects (i.e. as explanatory variables to quantify an average effect, *β*) in the mixed model. In a second step, we tested if *B. burgdorferi* s.l. or *B. afzelii* prevalence was higher at the range margin of *I. ricinus* (i.e. at high elevations). To this end, we compared the *B. burgdorferi* s.l. or *B. afzelii* prevalence in *I. ricinus* at the range margin (sites >1400 m.a.s.l.) to *B. burgdorferi* s.l. or *B. afzelii prevalence* in the core range of *I. ricinus* (sites < 1150 m.a.s.l.) by including range type (range core (C) vs range margin (M)) of each site as a fixed effect.

Finally, we focused on the patterns within the core range of *I. ricinus* by investigating the association between *B. burgdorferi* s.l. or *B. afzelii* prevalence and elevation, and the relationship between *B. burgdorferi* s.l. or *B. afzelii* prevalence and questing *I. ricinus* abundance within the core range of *I. ricinus*. The effect of questing *I. ricinus* abundance was investigated by including questing *I. ricinus* abundance and questing *I. ricinus* abundance^2^ as fixed effects.

To ensure independence between main and quadratic terms, all continuous variables were standardized (by subtracting the population mean from each sample and dividing it by the standard deviation). The significance of explanatory variables was assessed using likelihood ratio tests. The significance of the linear terms was assessed after removing the non-significant quadratic terms. Statistical analyses were performed using the package glmmADMB (Fournier et al., 2012; Skaug et al., 2014) in R 3.2.3 (R Core Team, 2013).

## 3. Results

Overall, 24.6% of *I. ricinus* nymphs (101 out of 411) were infected with *B. burgdorferi* s.l.. 83.2% of these infections were caused by *B. afzelii* (Table 1). Prevalence among sites varied between 0 and 47.1% for *B. burgdorferi* s.l., and 0 and 44.1% for *B. afzelii* (Table 1).

*Borrelia burgdorferi* s.l. prevalence decreased linearly with increasing elevation (Elevation^2^: *χ*^2^_1_ = 3.23, *P* = 0.072; Elevation: *χ*^2^ = 6.49, P = 0.011, *β* = −0.664, Fig.1B), whereas for *B. afzelii* the linear decrease with increasing elevation was marginally non-significant (Elevation^2^: *χ*^2^_1_ = 3.31, P = 0.069; Elevation: *χ*^2^_1_ = 3.46, *P* = 0.063; Fig.1C). *Borrelia* prevalence was 12.6 and 9.8 times higher in *I. ricinus* nymphs collected within the core range than at the range margin for *B. burgdorferi* s.l. (*χ*^2^_1_ = 9.55, *P* = 0.002, Fig.1B) and *B. afzelii* (*χ*^2^_1_ = 7.56, *P* = 0.006, Fig.1C), respectively.

Within the core range of *I. ricinus, Borrelia* prevalence was not significantly associated with elevation (*B. burgdorferi* s.l.: Elevation^2^: *χ*^2^_1_ = 0.02, *P* = 0.877; Elevation: *χ*^2^_1_ = 0.86, *P* = 0.353; *B. afzelii*: Elevation^2^: *χ*^2^_1_ = 0.06, P = 0.803; Elevation: *χ*^2^_1_ = 0.06, P = 0.800). Moreover, within the core range of *I. ricinus* no association between *Borrelia* prevalence and questing *I. ricinus* abundance was found for *B. burgdorferi* s.l. (Questing tick abundance^2^: *χ*^2^ = 2.09, *P* = 0.148; Questing tick abundance: *χ*^2^_1_ = 0.31, *P* = 0.576, Fig.1B) or *B. afzelii* (Questing tick abundance^2^: *χ*^2^_1_ = 2.00, *P* = 0.157; Questing tick abundance: *χ*^2^_1_ = 0.12, *P* = 0.733, Fig.1C).

## 4. Discussion

Laboratory experiments have shown that *B. burgdorferi* s.l. can alter the abiotic tolerance of *Ixodes* spp. (Herrmann and Gern, 2015, and references therein), suggesting that the spirochete may facilitate *Ixodes* spp. range expansion to marginal habitats, such as higher elevations or higher latitudes. Here we used a correlational approach to investigate whether patterns of *B. burgdorferi* s.l. prevalence in questing *I. ricinus* nymphs, and its variation with elevation, provide support for this hypothesis.

Overall, questing *I. ricinus* abundance was high and *B. burgdorferi* s.l. infection common in the study area: *I. ricinus* were found at 14 out of 15 sites, and *B. burgdorferi* s.l. infection in questing *I. ricinus* nymphs was detected at 12 sites. Above 1400 m.a.s.l., however, *I. ricinus* abundance strongly decreased and no *I. ricinus* were found above 1700 m.a.s.l.. *B. burgdorferi* s.l. prevalence in questing *I. ricinus* nymphs across sites was similar to previous reports from central Europe (e.g. Moran Cadenas et al., 2007, and references therein, Herrmann et al., 2013), but variation across sites was high (0 - 47% compared to 1 - 20% across Europe according to Rauter and Hartung, 2005), and it exceeded 40% at two sites of contrasting elevations (630 and 1123 m.a.s.l.).

As expected, the most common *B. burgdorferi* s.l. genospecies was *B. afzelii*, accounting for 83.2% of *I. ricinus* infections in our study. This is higher than what has been reported previously (7-68% of *I. ricinus* infections caused by *B. afzelii* across Europe; Rauter and Hartung, 2005). Differences across studies and locations are likely due to differences in the composition of the local host community, i.e. the relative abundance of rodent, bird and reptile hosts. Indeed the different *B. burgdorferi* s.l. genospecies are associated with different reservoir hosts. For example, *B. afzelii* and *B. bavariensis* are rodent specialists, *B. garinii* and *B. valaisiana* are bird specialists, and *B. burgdorferi* s.s. is a generalist that can infect birds and rodents (reviewed in Margos et al., 2011). Alternatively, strong bottlenecks within sites could lead to the local dominance of specific genospecies that may vary over time (e.g. Bruyndonckx et al., 2009; Criscione and Blouin, 2006).

We found no evidence that *B. burgdorferi* s.l. prevalence is higher at the range margins of *I. ricinus* (i.e. at higher elevations), indicating that the modification of *I. ricinus* physiology and behaviour by *B. burgdorferi* s.l. observed in the laboratory (Herrmann and Gern, 2015) plays a minor role in the ongoing colonisation process of marginal habitats in the wild. Rather, *I. ricinus* nymphs at the range margin had substantially lower *B. burgdorferi* s.l. prevalence.

This pattern may be due to different non-exclusive processes. The colder microclimate at high elevations may decrease the survival of *B. burgdorferi* s.l. in *I. ricinus* and slow down its multiplication and transmission. Effects of meteorological temperature on pathogen development and transmission efficiency (i.e. the development from the ingested infectious stage to the stage in the salivary glands) have been observed in malaria-mosquito (Eling et al., 2001; Noden et al., 1995) and trypanosome-tsetse fly systems (references in Moore et al., 2012). Although protozoan development is not directly comparable to bacterial multiplication, many bacteria similarly utilize their environmental temperature as a signal to determine their location and to regulate expression of a large set of proteins necessary for survival, multiplication, migration and transmission (Miller, 1989; Steinmann and Dersch, 2013). For example, the expressions of Outer Surface Protein A and C of *B. burgdorferi s.l.,* which are involved in its survival in the tick midgut and its dissemination into the tick salivary glands or into its vertebrate host, appear to be mediated by differences in ambient temperature (Schwan and Piesman, 2002). To our knowledge, no study evaluated the effect of meteorological temperature on the expression of vector-specific or host-specific proteins, or the expression of virulence genes in *B. burgdorferi* s.l..

Although temperature and other abiotic factors are often involved to explain the lower pathogen prevalence at higher elevations (Lafferty, 2009), the pattern may also be affected by other processes such as changes in host community, species abundance, habitat quality or population structure of hosts/vectors in marginal habitats. Indeed, Patot et al. (2010) found no effect of temperature on filamentous virus transmission efficiency, but observed a clear relationship between virus prevalence and the density of its host, a parasitoid wasp. Similarly, *B. burgdorferi* s.l. transmission may be lower in *I. ricinus* populations at the range margin that are more fragmented (Hanski, 1999), less dense (Patot et al., 2010) or/and less genetically diverse (Sexton et al., 2009) than populations at the core range. Similar processes may act on the reservoir host populations of *B. burgdorferi* s.l. (e.g. rodents and birds). However detailed knowledge about the composition of the host community, host abundance, *I. ricinus* infestation of these host populations as well as *B. burgdorferi* s.l. prevalence would be necessary to link host abundance and *B. burgdorferi* s.l. prevalence in questing *I. ricinus*. Furthermore, because edge populations are usually founded by only few individuals, stochastic processes may lead to the loss of pathogens. Thus, the combination of stochastic events and host and vector population structure that hinder *B. burgdorferi* s.l. transmission may lead to local extinctions of *B. burgdorferi* s.l. at the range edges (e.g. Phillips et al., 2010).

Our study is correlational, and thus has limitations. First, a time-based approach (which is less reliable than an area-based approach) was used to estimate questing tick abundance because of the rugged mountain terrain of the study area. Second, vegetation structure varies substantially across the elevation range and could affect tick detection despite a consistent sampling effort. However, the decrease of *I. ricinus* abundance above 1100 m.a.s.l. was observed in different life-stages having different questing behaviours and biological needs (Lemoine pers. obs.). Therefore, we are confident that the decrease of *I. ricinus* abundance is not a methodological artefact but describes an ecological pattern. Third, our approach does not allow to disentangle between expanding ticks bringing the pathogen with them (i.e., dispersal to a novel habitat) and the enzootic cycle being less likely to become established (i.e., establishment in a novel habitat, for example infecting less dense host populations). Fourth, whereas numerous studies in Europe and North America have documented a range expansion of ticks to higher elevations and latitudes due to climate change (reviewed in Medlock et al., 2013), we do not have the long-term monitoring data to directly demonstrate this range expansion at our study sites. Finally, the presence of questing *I. ricinus* ticks in the vegetation does not prove that *I. ricinus* populations are established locally, or will survive through the winter. High elevation habitats might thus represent a ‘ragged edge’ rather than an expansion front. Ultimately, an experimental approach, in which the *B. burgdorferi* s.l. infection status of *I. ricinus* is manipulated and the colonisation of marginal habitats is monitored would be required to test conclusively whether *B. burgdorferi* s.l. facilitates tick range expansion to marginal habitats in the wild.

In conclusion, in our correlative study we found no evidence that *B. burgdorferi* s.l.- induced changes in *I. ricinus* behaviour or physiology that facilitate *I. ricinus* range expansion to higher elevations in the Alps. Rather, questing *I. ricinus* in marginal habitats are less likely to carry *B. burgdorferi* s.l.. These findings show that when bitten by a tick, the risk of human *B. burgdorferi* s.l. infection is lower, rather than higher, in regions where *I. ricinus* is newly emerging due to climate change. Low *B. burgdorferi* s.l. prevalence at *I. ricinus* range margins may enhance population growth and competitive ability of hosts and vectors. Less infected, hosts may, for example, invest differently in immunity and reproduction than hosts in core populations (White and Perkins, 2012), which can affect host-parasite interactions when the parasite finally invades host populations at the range margins (Kelehear et al., 2012). A better understanding of eco-evolutionary processes between pathogens, vectors and hosts at range margins, and their effect on pathogen life-history and virulence evolution, will therefore be a fruitful next step (e.g. Kelehear et al., 2012), and will contribute to a better prediction of zoonotic disease risks in regions where vectors and pathogens are newly emerging due to climate change.

## Acknowledgements

The study was financially supported by the University of Zurich Research Priority Program ‘Evolution in Action: From Genomes to Ecosystems’ (to ML and BT), the Swiss National Science Foundation (PMPDP3_151361 and PMPDP3_161858 to ML; PP00P3_128386 and PP00P3_157455 to BT), the Baugarten Stiftung & Stiftung für wissenschaftliche Forschung an der Universität Zürich (STWF-17-027), and the Georges und Antoine Claraz-Schenkung. We thank the local authorities for their permission to collect ticks in their municipal areas, the field assistants for their help with data collection, Reto Lienhard, ADMED Microbiologie, for providing the reference strains, and Kay Lucek and three anonymous reviewers for helpful comments on the manuscript.

Declarations of interest: none

